# Membrane-associated polymerases deliver most of the actin subunits to a lamellipodial network

**DOI:** 10.1101/2025.03.24.645090

**Authors:** Kristen Skruber, David Sept, R. Dyche Mullins

**Affiliations:** Department of Cellular and Molecular Pharmacology Howard Hughes Medical Institute UCSF School of Medicine San Francisco, CA 94143; Department of Biomedical Engineering University of Michigan Ann Arbor, MI

## Abstract

Actin filaments are two-stranded protein polymers that form the basic structural unit of the eukaryotic actin cytoskeleton. While filaments assembled from purified actin *in vitro* elongate when soluble monomers bind to free filament ends, in cells the mechanism of filament elongation is less clear. Most monomeric actin in the cytoplasm is bound to the accessory protein profilin, and many regulators of filament assembly recruit actin-profilin complexes to membrane surfaces where they locally accelerate filament elongation. Employing quantitative live-cell imaging of actin-profilin fusion proteins and biochemically defined mutants of the branched actin regulator, WAVE1, we find that only ∼25% of the actin in leading-edge lamellipodial networks enters directly from solution, while the majority enters via membrane-associated polymerases.

## Introduction

The goal of this study is to determine the primary pathway by which actin monomers become subunits of branched filament networks in living cells. Specifically, we aim to measure the relative contributions of two alternate pathways: (1) direct filament elongation from soluble monomers in the cytoplasm^1^ versus (2) indirect elongation via membrane-associated polymerases^2^. Filament elongation by soluble monomers is a fast, diffusion-limited reaction^1^ that is well described by a set of first- and second-order rate constants^3^. Although elongation by membrane-associated polymerases requires two steps—loading actin monomers onto the membrane and then transferring them to nearby filament ends—in vitro this process can be much faster than direct incorporation from solution when the surface density is sufficiently high^4^ (Bieling, 2018). The rates of direct and membrane-mediated filament elongation, however, depend on several factors, including the concentration of soluble actin, the density of polymerase molecules on the membrane, load forces, and the proximity of filaments to the membrane surface^5^. For this reason, the relative contributions of the two pathways to filament formation in cells remain unknown.

Self-assembly of branched actin networks created by the Arp2/3 complex generates force to drive membrane movement, including in lamellipod protrusion^6^, endocytosis^7^, phagocytosis^8^, cell-cell adhesion^9^, cell fusion^10^, plasma membrane repair^11^, and autophagosome formation^12^. In each of these processes, upstream signaling molecules recruit regulators of actin filament assembly, called nucleation promoting factors (NPFs)^13^, to a membrane surface. When activated on the membrane, these factors, generally members of the WAVE/WASP protein family, create a branched filament network by locally stimulating the Arp2/3 complex to autocatalytically create new filaments that grow from the sides of preexisting filaments^14,15^.

In addition to promoting Arp2/3-dependent filament nucleation, WAVE/WASP-family proteins also bind actin monomers and actin-profilin complexes^16^. When clustered on a surface, these binding sites work together to create an efficient, distributive actin polymerase that accelerates elongation of nearby filaments^4^. The magnitude of this acceleration, relative to the direct incorporation of soluble monomers, determines the fraction of total actin that is delivered to the network by these surface polymerases^5^.

### Probes for tracking the route of actin from the cytoplasm into a filament network

Because most soluble actin monomers in the cell are bound to profilin^17^ we focused on the polymerase activity of the proline-rich profilin-binding domains found in nucleation promoting factors and other membrane-associated actin regulators, such as ENA/VASP-family proteins^18^ and formins^19^. Wildtype actin-profilin complexes can elongate filaments *both* from solution^20^ *and* via surface-associated polymerases^4^. Complexes containing mutant profilins defective for binding polymerase proteins, such as H113S (hereafter referred to as proline-binding deficient, PBD)^21^, elongate filaments *only* via the solution pathway. Comparing the rates at which wildtype and mutant profilins support network growth would, therefore, reveal the relative contributions of solution and surface-mediated pathways (Fig. 1A). Unfortunately, profilin dissociates rapidly from the end of a growing filament^17,20^ and thus does not ‘label’ the filaments it helps construct.

**Figure 1.**
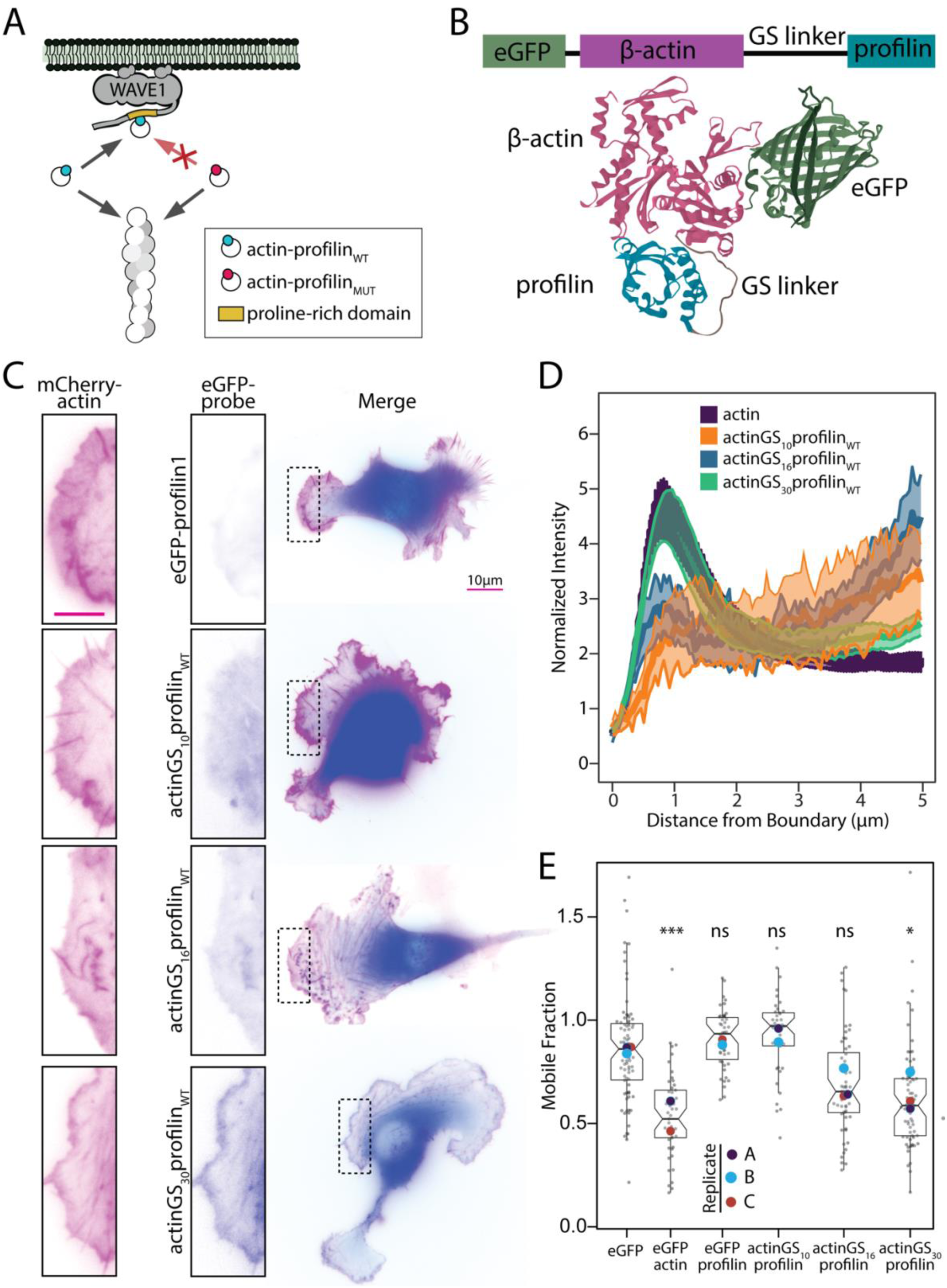
Tethered actin profilin constructs incorporate into actin networks. A. Schematic representation of the route actin takes into filaments through the actin-binding protein profilin. The covalently linked actin-profilin probe can enter filaments directly from the cytosol and by first binding to the proline-rich region of NPFs like WAVE1. A similar probe that contains a modified profilin that can still incorporate into filaments but cannot bind proline-rich domains (actin-profilin_MUT_) only enters the filament directly from the cytosol. B. Illustration of actin-profilin orientations for linked constructs using previously published structures (PDB:2BTF) with either 10, 16 or 30 alternating glycine-serine residues between the N terminus of β-actin and the C-terminus of profilin. EGFP is linked to the N terminus of β-actin for probe visualization. C. Fluorescence of B16F1 cells transiently expressing indicated constructs. Scale bar represents 5µm. D. Averaged linescans at the leading edge of cells expressing actin-profilin probes with different linker lengths, where the cell edge = 0. The bands depict 95% confidence intervals. For all conditions, n = 100 lines drawn from 10 cells. Lines are normalized to the cell mean intensity. Only the probe with the longest linker shows a similar increase at the leading edge to eGFP-actin. E. Quantification of mobile fraction of recovery curve after photobleaching per cell indicating that both actin and actinGS_30_profilin readily encorporate into filaments, as they both exhibit substantial nonmobile fractions. Number of cells per experiment per round for both A and B are as follows with cell number indicated in parenthesis: eGFP (37,23,22) eGFP-actin_WT_ (22,27), eGFP-profilin1 (11,9,26), β-actinGS_10_profilin (23,18), β-actinGS_16_profilin (21,12,23), β-actinGS_30_profilin (16,35,14). One-way ANOVA and Tukey’s multiple comparison tests are comparisons with eGFP.

To compare the abilities of wildtype and mutant profilins to support filament elongation, we made use of an observation by Carlier and co-workers that growing actin filaments can incorporate covalently crosslinked actin-profilin complexes at almost the same rate as normal, non-crosslinked complexes^22^. Instead of chemically crosslinking the two proteins, however, we created genetically encoded actin-profilin (AxP) complexes by attaching the N-terminus of a profilin isoform (either profilin1 or profilin2) to the C-terminus of β-actin, via a flexible Gly-Ser-rich linker (Fig. 1B). We reasoned that, when an AxP complex binds a growing filament end, the profilin component could disengage from its actin partner while still remaining tethered to the filament. This tethering would provide the filament with a ‘memory’ of the profilins that helped construct it. To follow these probes in live cells, we fused a fluorescent protein (enhanced Green Fluorescent Protein, eGFP) to the N-terminus of the actin (Fig. 1B).

The polymerizability of covalently linked actin-profilin complexes depends critically on the geometry of their linkage^22,23^. We created and tested fusion constructs of β-actin and profilin linked by repeating Gly-Gly-Gly-Ser sequences of different lengths: 10, 16, or 30 amino acids (Fig. 1B). We tested the relative competence of these variable-linker AxP probes to enter branched actin networks by monitoring their dynamics in lamellipodia of spreading B16F1 mouse melanoma cells. The leading edge of these cells is a well characterized model system for studying the assembly of branched actin networks^24^.

Lamellipodia of spreading B16F1 cells comprise thin, convex protrusions, closely apposed to the substrate and supported by a dynamic actin network. In cells expressing mCherry-actin, this lamellipodial actin network appears as a broad peak in fluorescence intensity near the cell edge (Fig. 1C, left column). We compared the localization and mobility of eGFP-labeled AxP probes to a co-expressed mCherry-actin standard using two assays in live cells: line-scans of normalized fluorescence intensity across the cell edge and fluorescence recovery after photobleaching (FRAP) near the cell edge (Fig. 1). We also employed eGFP-actin as a positive control for network incorporation and both eGFP and eGFP-profilin1 as negative controls. Line scans of fluorescence intensity reveal the relative accumulation of the probe along a single slice through the leading edge (Fig. 1C-D). Fluorescence recovery reveals the fractions of monomeric and filamentous actin at a given point in the network: the rapidly recovering fraction of the signal reflects highly mobile monomers^25^ while the slowly recovering fraction reflects less mobile filaments^26^.

By both assays, the polymerization competence of an AxP probe increases with linker length. Specifically, the probe with the shortest, 10-amino acid, linker (actinGS_10_profilin_WT_) failed to incorporate to detectable levels in at the leading edge (Fig. 1C-D) and, in FRAP assays, exhibited a mobile fraction indistinguishable from both eGFP and eGFP-profilin1 (Fig. 1E, Fig. S1). The 16-amino acid linker (actinGS_16_profilin_WT_) supported partial network incorporation compared to mCherry-actin in the two assays (Fig. 1C-E). Remarkably, the probe with the 30 amino acid linker (actinGS_30_profilin_WT_) was indistinguishable from the actin standard in both leading edge line scans (Fig. 1D) and fluorescence recovery assays (Fig. 1E), indicating that the polymerization competence of this GFP-actin-profilin complex is approximately equal to that of mCherry-actin.

Why do longer linkers enable more rapid filament assembly? Based on previous studies of Gly-Ser-containing peptides^27^, we estimate equilibrium lengths of 21, 27, and 37 Å for the 10-, 16-, and 30-amino acid linkers respectively. In solution, these flexible polypeptides behave as entropic springs, capable of exerting a pulling force when their end-to-end distance is held above the equilibrium value. In an actin-profilin complex, the distance from the C-terminus of actin to the N-terminus of profilin is 31.2 A^28^, slightly shorter than the estimated end-to-end distance of the 30-amino acid linker. This suggests that the shorter linkers may generate tension between the linked proteins, possibly perturbing their structures. To test this idea, we compared molecular dynamics simulations of the untethered actin-profilin complex with those of AxP constructs containing 10-and 16-amino acid linkers (Fig. S3). By principal component analysis, the unlinked complex and the complex with the 16-amino acid linker have almost indistinguishable conformations, while the 10-amino acid linker induces a significant structural perturbation.

Finally, to test whether the profilin-actin interface is required for polymerization of the covalently linked complexes we linked actin to a profilin1 mutant (R88E) that is defective in binding actin (actinGS_30_profilin_R88E_)^29^. By fluorescence microscopy and FRAP analysis, this AxP construct failed to accumulate in lamellipodial networks as did a non-polymerizable mutant actin (Fig. 2A-B). This result argues that the actin and profilin components of a functional AxP probe act together as a unit, mimicking the behavior of a native actin-profilin complex.

**Figure 2.**
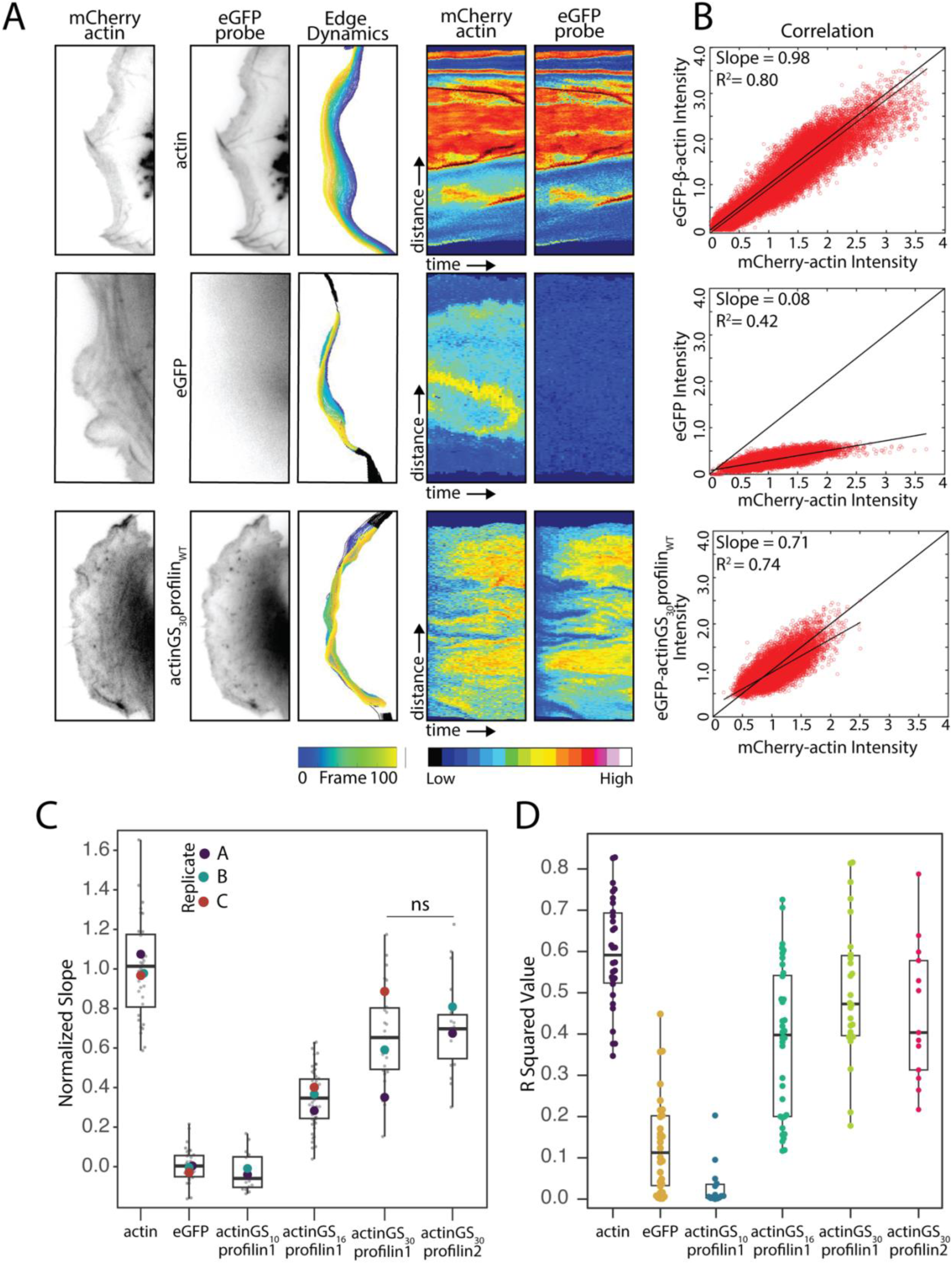
β-actin-profilin construct intensity correlates with intensity of β-actin in cells. A. From left to right: Representative images of a dynamic leading. The β-actin channel was used to identify the leading edge. Edge dynamics show the automated segmention of the leading edge over consecutive frames. Adaptive kymographs remap fluorescence intensity at the leading-edge position to the y-axis and time to the x-axis to visualize dynamic incorporation of linked constructs at the leading edge. Color map: blue (low fluorescence intensity) to red (high fluorescence intensity). Scale bar represents 5µm. B. Example correlation plots between normalized intensity values for each probe co-expressed with actin. All correlation lines are normalized to the average correlation for actin with itself, displayed as a thicker black line through the origin. For correlation with actin channel, measurement is between eGFP on the relevant protein (β-actin, eGFP alone, or actinGS_30_profilin probe) and mCherry β-actin. (C-D) Quantified slopes and R squared values for correlation plots between each probe with the β-actin channel. Number of cells per experiment per round are as follows: actin (12,11,14), eGFP (28,19,11), actinGS_10_profilin1_WT_(18,19,14), actinGS_16_profilin1_WT_ (23,17,18), actinGS_30_profilin1_WT_ (15,7,26), actinGS_30_profilin2_WT_ (25,7,25). Replicate means in C are represented by large dots superimposed on the cell-level data. One-way ANOVA and Tukey’s multiple comparison tests were performed in C on the mean of each replicate.

### Quantifying relative rates of AxP probe incorporation into lamellipodial networks

Line scans and fluorescence recovery assays provide localized snapshots of labeled actin assembly but do not quantify network incorporation rates. To more accurately determine the relative rates at which AxP probes incorporate into actin networks, we employed automated edge detection to measure and correlate leading-edge actin assembly across space and time^30^. Briefly, we performed time-lapse fluorescence microscopy on cells co-expressing an eGFP-labeled AxP probe together with mCherry-actin. We collected simultaneous imaging pairs of both fluorophores (e.g. Fig. 2A, left two columns) at 1-second intervals for 100 seconds. We then used an adaptive space-time correlation method to measure fluorescence intensity in both channels along the same moving cell edge in every image of the time-lapse sequence (Fig. 2A, third column). After normalizing the raw intensities to the total fluorescence of each probe in the cell, we generated two-dimensional space-time plots (a.k.a. kymographs) of relative intensity across both the length of the cell edge and the duration of the time-lapse sequence (Fig. 2A, columns 4-5). To quantify incorporation rate of the AxP probe relative to the mCherry-actin reference, we plotted the intensity of probe versus reference fluorescence at every point of the kymograph (Fig. 2B). We then used linear regression to determine a best-fit slope and R^2^ value for this plot. We interpret the slope as the rate of probe incorporation relative to the actin standard and the R^2^ value as a measure of whether fluctuations in the two intensities arise from the same underlying process. To validate this approach, we compared eGFP-actin to the mCherry-actin reference and obtained an average slope of 0.98 ± 0.07 with an R^2^ value of 0.6 ± 0.05 (Table 1 ; 32 cells from three biological replicates). We also determined a zero-incorporation baseline slope for our experiments of 0.10 ± 0.03, by computing the correlation of soluble eGFP and mCherry-actin (Table 1; 35 cells from three replicates). Soluble eGFP does not incorporate into branched actin networks, and so the residual, non-zero slope of this plot likely reflects small changes in optical path length that are weakly correlated with the density of filamentous actin in the network. The low R^2^ value (0.13 ± 0.05) for the linear fit indicates that most of the eGFP fluorescence intensity is better explained by random fluctuations than by actin-dependent changes in the signal.

**Table 1.** Slopes and linear correlation coefficient for correlation of indicated probe with actin. The normalized slope (represented as % Slope_NORM_) normalizes each value to eGFP and β-actin as 0 and 100% incorporation, respectively. Error is reported with a 95% confidence interval.

| Correlation with<br>β-Actin | Slope | % Slope<sub>NORM</sub> | R<sup>2</sup> |
| --- | --- | --- | --- |
| eGFP | $0.10 \pm 0.03$ | 0 | $0.13 \pm 0.04$ |
| β-Actin<sub>WT</sub> | $0.94 \pm 0.08$ | 100 | $0.60 \pm 0.05$ |
| ActinGS<sub>10</sub>PFN1<sub>WT</sub> | $0.09 \pm 0.04$ | 0 | $0.03 \pm 0.03$ |
| ActinGS<sub>10</sub>PFN1<sub>PBD</sub> | $0.09 \pm 0.04$ | 0 | $0.05 \pm 0.02$ |
| ActinGS<sub>16</sub>PFN1<sub>WT</sub> | $0.39 \pm 0.04$ | 34 | $0.38 \pm 0.06$ |
| ActinGS<sub>16</sub>PFN1<sub>PBD</sub> | $0.25 \pm 0.05$ | 17 | $0.34 \pm 0.10$ |
| ActinGS<sub>30</sub>PFN1<sub>WT</sub> | $0.69 \pm 0.09$ | 70 | $0.50 \pm 0.06$ |
| ActinGS<sub>30</sub>PFN1<sub>PBD</sub> | $0.36 \pm 0.03$ | 17 | $0.37 \pm 0.05$ |
| ActinGS<sub>30</sub>PFN2<sub>WT</sub> | $0.70 \pm 0.10$ | 71 | $0.45 \pm 0.10$ |

We quantified the relative rate of incorporation (r) of each AxP probe with the formula,

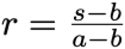

where s is the average slope of the space-time correlation plot of the probe; b is the baseline slope of soluble eGFP; and a is the slope of the eGFP-actin positive control. The relative rates of incorporation for profilin1-containing AxP probes made with 10-, 16-, and 30-amino acid linkers are 0, 0.34, and 0.7, compared to the mCherry-β-actin standard (Table 1, column 3). The low correlation slope (0.09 ± 0.04) and low R^2^ value (0.03 ± 0.03) of the AxP construct with a 10-amino acid linker (actinGS_10_profilin_WT_) suggest that this complex is completely incompetent for polymerization at the leading edge. To determine whether different profilin isoforms promote different rates of filament growth, we also compared AxP constructs made with profilin1 and profilin2, the two major forms of profilin found in mammalian cells^31^. When connected by a 30-amino acid linker, both profilin isoforms promoted the same relative rate of network incorporation: 0.70 ± 0.11 for profilin1 versus 0.71 ± 0.10 for profilin2 (Table 1). In addition, the R^2^ values for the correlation plots of the two probes were similar to that of the eGFP-actin control (Table 1).

### Profilin-poly-proline interactions mediate most filament assembly from profilin-actin

After constructing and validating the AxP probes, we next used them to measure the fraction of actin that enters a lamellipodial network via interaction between profilin and the proline-rich binding sites on membrane-associated polymerases. Specifically, we compared the relative incorporation of AxP probes containing wildtype profilin (i.e. actinGS_30_profilin_WT_) to those with mutated polymerase binding sites (i.e. actinGS_30_profilin_PBD_). We first performed fluorescence ratio imaging, in which we divided the normalized intensity of wildtype and mutant AxP probes at every point in the image by the intensity of co-expressed mCherry-actin (Fig. 3A). These images revealed a significant defect in leading edge accumulation of the mutant probe (actinGS_30_profilin_PBD_) compared to the wildtype (actinGS_30_profilin_WT_). Linescans across leading-edge lamellipodia revealed a similar defect in actinGS_30_profilin_PBD_, which peaked at a value less than 30% of the wildtype probe, actinGS_30_profilin_WT_ (Fig. 3B). We also observed a significant drop in leading-edge fluorescence intensity of the mutant, 16-amino-acid-linker construct, actinGS_16_-profilin_PBD_ compared to wildtype, actinGS_16_-profilin_WT_ (Fig. S4). Finally, we used space-time correlation along dynamic lamellipodia to quantify the incorporation defect caused by mutating the polymerase binding site of profilin. For profilin1-containing probes the relative incorporation rate (compared to mCherry-actin) decreased from 0.70 ± 0.11, for actinGS_30_profilin_WT_, to 0.17 ± 0.02, for actinGS_30_profilin_PBD_ (Fig. 3C-E, Table 1).

**Figure 3.**
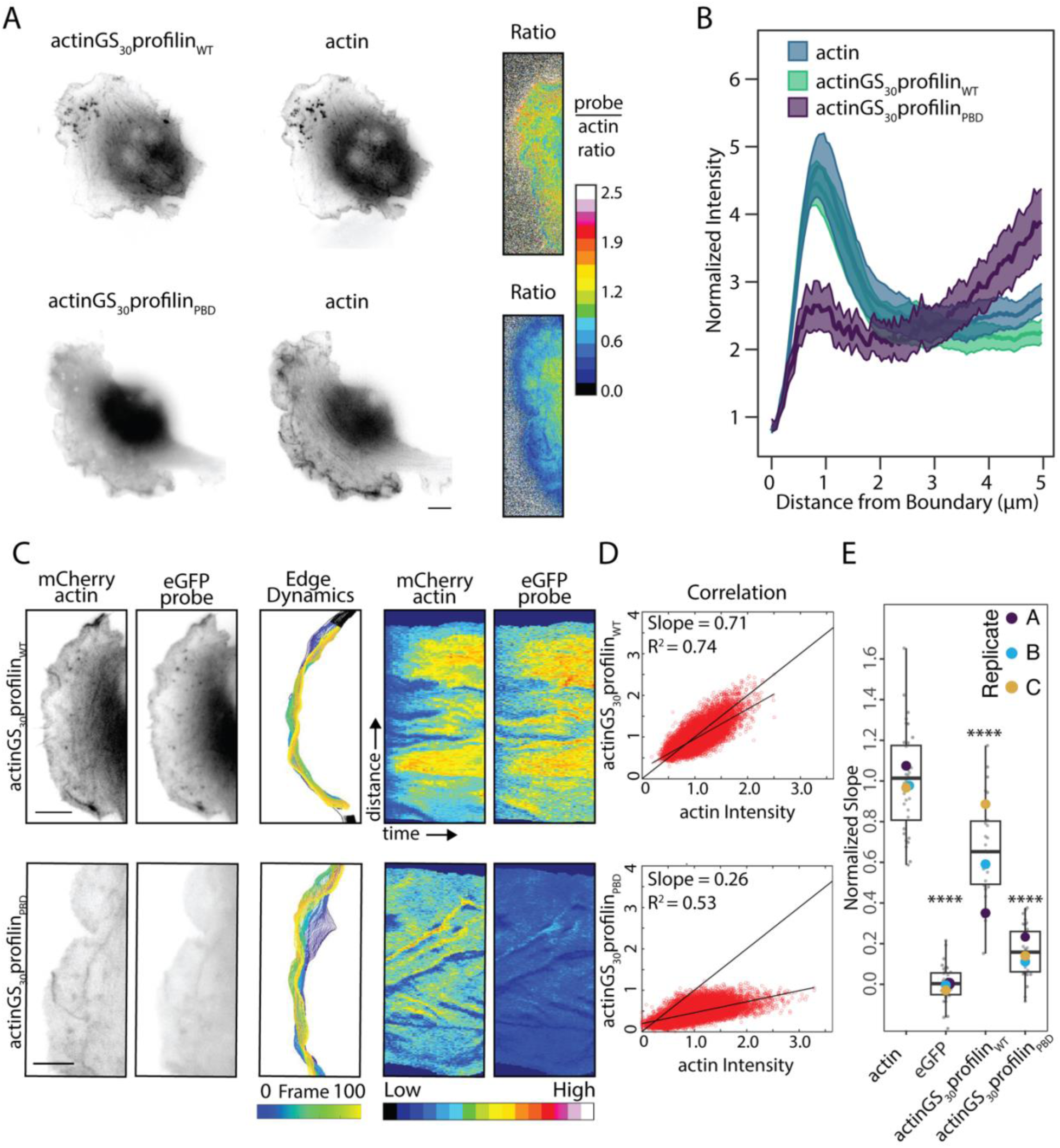
Profilin’s ability to bind Proline-Rich Domains is crucial for localization of actin to the leading edge. A. Representative images displaying localization to the leading edge of actinGS_30_profilin1_WT_ or actinGS_30_profilin1_PBD_ (proline-binding deficient that cannot bind NPFs). Ratiometric images on the right display ratio images generated by dividing actinGS_30_profilin_WT_ (top) or actinGS_30_profilin_PBD_ (bottom) channel by the β-actin channel (higher value indicates more colocalization of the probe with actin), which indicates that the actinGS_30_profilin_PBD_ probe fails to colocalize with actin. Scale bar represents 5µm for inset and 10µm for whole image. B. Averaged line scans of actinGS_30_profilin_WT_ and actinGS_30_profilin_PBD_ fluorescence intensity at the leading edge of cells. Only the actinGS_30_profilin_WT_ probe accumulates at the leading edge like actin. Distance indictates distance from the cell edge where the edge = 0. The transparent bands depict 95% confidence intervals. For all conditions, n = 200 lines drawn from 20 cells. Lines are normalized to the cell mean intensity. C. From left to right: Representative images of a dynamic leading edge. The actin channel was used to identify the leading edge. Scale bar represents 5µm. Edge dynamics and adaptive kymographs as described in Figure 2. Color map: blue (low fluorescence intensity) to red (high fluorescence intensity). D. Example correlation plots between normalized intensity values for each probe co-expressed with β-actin, as described in Figure 2. E. Quantified slopes for correlation plots between each probe with the corresponding actin channel. Number of cells per experiment per round are as follows: actin (12,11,14), eGFP (28,19,11), actinGS_30_profilin1_WT_ (15,7,26), actinGS_30_profilin2_WT_ (25,7,25). Replicate means are represented by large dots superimposed on the cell-level data. Number of cells per experiment per round are as follows: actin (12,11,14), eGFP (28,19,11), β-actinGS_10_profilin1_WT_ (18,19,14), β-actinGS_30_profilin1_WT_ (15,7,26), β-actinGS_30_profilin1_PBD_ (10,22,17). Replicate means are represented by large dots superimposed on cell-level data. One-way ANOVA and Tukey’s multiple comparison tests were performed in E. on unnormalized replicate means.

We attribute lower incorporation of the mutant AxP probes to decreased binding of membrane-associated (as opposed to soluble) polymerases, in part because most proline-rich profilin binding partners, including WAVE/WASP, ENA/VASP, and formin-family proteins, are active when localized to membrane surfaces. More specifically, the profilin-binding domains of the WAVE/WASP family proteins that create lamellipodia act as polymerases *only* when clustered on a two-dimensional surface^4^. Given that this decrease reflects loss of the polymerase-mediated component of polymerization, the fraction of total actin provided to the network by surface polymerases (f_surf_) is given by

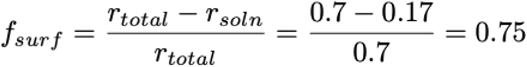

Where r_total_ and r_soln_ are the measured incorporation rates of wildtype and polymerase-binding-mutant AxP constructs (i.e. actinGS_30_profilin_WT_ and actinGS_30_profilin_PBD_). Thus, our results reveal that 75% of actin enters a lamellipodial network via interaction between profilin and membrane surface-associated polymerases.

To determine whether expression of covalently linked actin-profilin complexes perturbs actin dynamics at the leading edge, we measured the rate of retrograde flow of lamellipodial actin in the presence of each AxP construct. The baseline rate of retrograde flow in the absence of AxP constructs, based on kymographs of co-expressed mCherry-actin fluorescence, was 0.10 ± 0.006 µm/s (22 cells from 3 biological replicates). Expressing most AxP constructs had no effect on this rate. The only exception was the proline binding-deficient complex, actinGS_30_profilin_PBD_, which slightly decreased the rate of mCherry-actin retrograde flow (Fig. S5), consistent with a mildly deleterious effects on network growth.

Interestingly, AxP constructs made with wildtype profilin1 are sensitive to actin-profilin linker length, but constructs made with proline-binding deficient (H133S) profilin1 incorporate at the same low rate regardless of whether the linker is 16 or 30 amino acids long (Fig. 3C, Table 1). We observe a similar equivalence between the mobile fractions of complexes with 16- and 30-amino acid linkers (actinGS_16_profilin_PBD_ and actinGS_30_profilin_PBD_) in FRAP assays (Fig. S1). These results suggest that incorporation of crosslinked actin-profilin complexes from membrane-associated polymerases is more sensitive to linker length than incorporation of soluble complexes.

### Profilin-binding sites on WAVE1 promote lamellipodial actin network formation

Our results argue that profilin-binding polymerases mediate most actin assembly in a lamellipodial network, but which polymerases do this work? Leading edge membranes are packed with proline-rich domains from ENA/VASP proteins, formins, and multiple WAVE/WASP-family nucleation promoting factors^32^. Depleting cells of ENA/VASP proteins or individual formins can alter cellular morphology, but does not kill lamellipod formation^33,34^. In contrast, the nucleation promoting factors, WAVE1 and WAVE2, are required for branched actin network formation at the leading edge^35,36^. We previously found that the proline-rich region of WAVE1 is sufficient to create an effective polymerase when clustered on a surface *in vitro*, but is this activity important in live cells? Intriguingly, Buracco and co-workers recently demonstrated that WAVE1 truncation mutants that contain the proline-rich region and thus profilin binding sites, but lack the Arp2/3-activating sequences, rescue lamellipod formation in B16F1 cells^35^. These authors did not interpret their result in terms of profilin binding or polymerase activity but suggested that lamellipod rescue was due to interaction with other actin regulators that contain SH3 domains, such as Abi1.

To directly test the importance of profilin binding to WAVE1’s ability to form branched actin networks (Fig. 4A), we created WAVE1 constructs with point mutations in the three highest affinity profilin binding sites^4^ within the proline-rich region of the protein (WAVE1-PRD_MUT_). Importantly, these mutations (Fig. 4B) do not disrupt any known SH3 binding sites. We then compared the ability of wildtype WAVE1 versus the WAVE1-PRD_MUT_ mutants to rescue lamellipod formation in WAVE1/WAVE2 double knockout B16F1 cells^36^ (Fig. 4C-E). We assessed lamellipod architecture by two-channel, live-cell fluorescence microscopy of eGFP-labeled WAVE1 rescue constructs with mCherry-labeled F-tractin, as a probe for filamentous actin. The F-tractin peptide has a relatively low affinity for filamentous actin and labels branched actin networks better than other live cell actin probes, such as Lifeact or Utrophin^37^.

**Figure 4.**
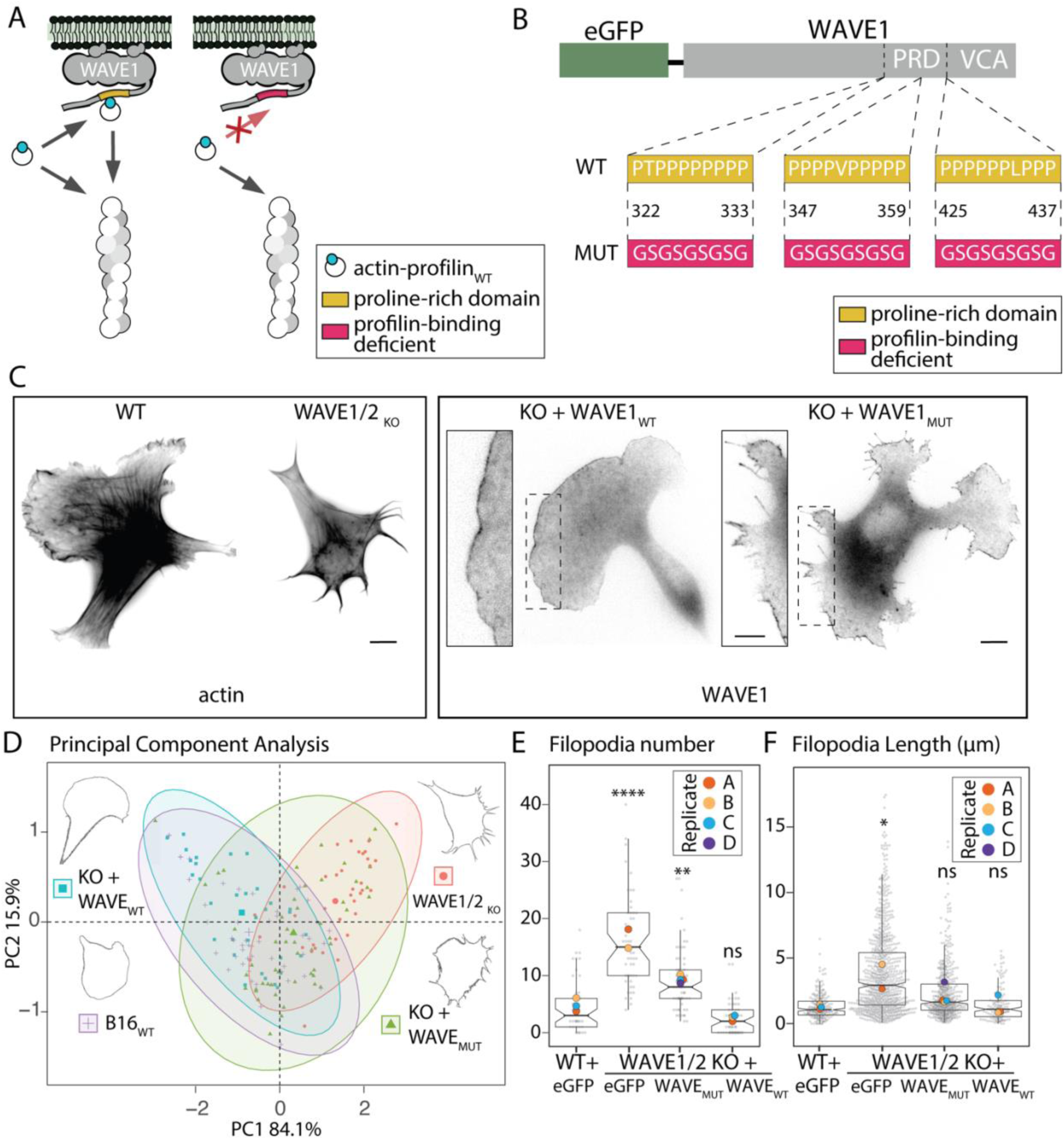
Effect of proline-rich domains on WAVE1 localization and generation of actin structures. A. Cartoon illustrating actin polymerization with the WAVE1 regulatory complex with the proline-rich region intact (left) and a WAVE1 with its proline-rich region mutated so that profilin cannot bind (right). B. Diagram of the PVCA domain of WAVE1 depicting modifications made to the proline-rich domain to alter binding of profilin-actin (profilin-binding deficient). C. Fluorescence of B16F1 depleted of both WAVE1 and 2 isoforms (WAVE1/2 double knockout) and rescues transiently expressing indicated constructs. B16F1 wild-type cells are shown for comparison. EGFP-WAVE1 with a modified proline-rich domain (WAVE1_MUT_) is shown in comparison to the intact domain (WAVE1_WT_), with magnified insets of the leading edge. The wildtype WAVE1 rescues the typical lamellipodia phenotype, while cells with the WAVE1_MUT_ that cannot bind profilin exhibit many filopodia, more similar to the double knockout cells. Scale bar represents 5 µm for inset, 10 µm for whole image. D. Principal component analysis of cell shape for various cell lines. The cells rescued with wild type WAVE1 appear more similar to wild type B16F1 cells, while cells transiently expressing only the WAVE1_MUT_ construct are intermediate between the wild type and double-knockout cells. The first principal component (PC1, x-axis) explains 84.1% of the variation in the data, while the second principal component (PC2, y-axis) represents 15.9% of the variation. Confidence ellipses are drawn at 95% for each group. Means for each group are displayed by larger sizes and representative outlines of cell shape for each mean are displayed in the corners. E. Filopodia number per cell. B16F1 wildtype cells (16, 14, 15), double-knockout cells (16, 37), and double-knockout cells transiently expressing either WAVE1_MUT_ (10,10,12,23) or WAVE1_WT_ (8,15,20). F. Filopodia length in microns, n= filopodia number per round and are as follows: (58,88,69), (290,536), (79,99,11), and (52,20,60), respectively. Replicate means are represented by large dots superimposed on cell-level data. One-way ANOVA and Tukey’s multiple comparison tests were performed on replicate means.

We find that WAVE1 mutants defective for profilin binding fail to fully restore lamellipod formation in WAVE1/2 double knockout B16F1 cells. Rather than a smooth lamellar protrusion, the leading edge of WAVE1_MUT_-expressing cells was also dominated by filopodia (Fig. 4C, Fig. S6), similar to the double knockout cells (Fig. S6). Using FiloQuant^38^ to automatically detect and measure filopodia, yielded an average number of 9.2 ±1.4 structures/cell (Fig. 4E) and an average length of 2.3 ± 0.2 µm (Fig. 4F). This was slightly lower than in WAVE1/2 depleted cells where the average length was 3.9 ±0.2 µm with an average of 15.8 ±2.1 filopodia/cell. WAVE1 expression with proline-rich regions intact in WAVE1/2 knockout cells decreased the filopodia length (1.5 ± 0.2 µm) to values indistinguishable from that of wildtype B16F1 cells (1.3 ±0.1 µm) with even fewer filopodia per cell than those with the proline-rich domain completely ablated (2.5 ±0.7 structures/cell), indicating restoration of the lamellipod.

More generally, B16F1 cells expressing the WAVE1_MUT_ defective for recruitment of profilin-actin were morphologically different from both WAVE1/2 knockout cells and cells expressing wildtype WAEV1. By principal component analysis of shape metrics, wildtype B16F1 cells were *least* similar to the WAVE1/2 double knockout cells and most similar to double knockouts expressing WAVE1_WT_ (Fig. 4D, Fig. S7). Cells expressing WAVE1-PRD_MUT_ displayed an intermediate phenotype but were closer to double knockout cells than to either wildtype or knockoutcells expressing WAVE1 in coordinate space. From these data, we conclude that the profilin binding proline-rich region of WAVE complexes on the membrane are instrumental in delivering actin to the barbed ends of growing actin fialments at the leading edge of cells.

## Discussion

Using profilin-linked actin, we were able to directly measure the fraction of monomers entering actin networks from a membrane surface in living cells. These results support the model (Mullins et al., 2018) that the majority of monomers (∼75%) incorporate into filaments from polymerases associated with the membrane surface. Revoking access to proline-rich sites reduces incorporation of actin-profilin, suggesting a hard limit (∼25%) on the fraction of monomers that can enter branched actin networks directly from the cytoplasm.

Previous *in vitro* experiments demonstrating that clustered profilin binding sites locally accelerate actin filament elongation were performed at non-physiologically low concentrations of soluble actin-profilin^4^. For this reason, it was formally possible that, at high cytoplasmic actin-profilin concentrations (100-150 µM in many cells), direct incorporation of soluble actin-profilin would overload the contribution from membrane-associated polymerases. Theoretical analysis suggests that at realistic leading-edge densities, polymerases could continue to deliver the majority of actin subunits to lamellipodial networks even at extremely high (supra-physiological) actin-profilin concentrations^5^. Our observation that 75% of actin-profilin is delivered to branched actin networks by membrane-associated polymerases is consistent with this theoretical analysis, given actin-profilin concentrations between 100 and 150 µM and membrane polymerase densities in the 2000-6000/µm^2^ range.

Our results reveal that, similar to intracellular signaling systems^39^, the actin cytoskeleton employs membrane localization and dimensionality reduction to accelerate critical protein-protein interactions^5^. This fact is important for several reasons. Firstly, it means that simple mass action cannot adequately describe actin filament elongation in cells and that numerical simulations of cellular actin assembly must employ more sophisticated models. Secondly, it explains the ubiquity of proline-rich profilin binding sites found in membrane-associated actin regulators across eukaryotic phyla and helps make sense of otherwise cryptic mutant phenotypes. Finally, nucleation promoting factors have not *two*, but *three* ways to control the architecture and function of a branched actin network. In addition to tuning the rate of filament nucleation^13,40^ and the extent of actin-membrane tethering^41^, WAVE/WASP family proteins can locally control the speed of filament elongation through direct delivery of actin subunits. The interplay between these three network-defining parameters is the key to understanding the biophysical mechanisms underlying the formation of branched actin networks in living cells.

## Methods

### Experimental Design and Statistical Analysis

Experiments, referred to graphically as “Replicate”, were performed three times, unless indicated otherwise. Data were visualized using R studio (Version 2022.07.1) and statistical analysis was performed using GraphPad Prism version (9.2.0, GraphPad Software, San Diego, CA) or R. Boxplots generated in R visualize seven summary statistics (one median, two notches, two hinges, two whiskers) at their default values. The notches are 1.58 times the interquartile range over the square root of the number of values, which is approximately a 95% confidence interval. The lower and upper hinges are the first and third quartiles, respectively. Whiskers are displayed as 1.5 times the interquartile range. For statistical analysis comparing three or more groups, one-way ANOVA and Tukey’s (or Dunn’s) multiple comparison tests were performed as indicated in the Figure legends. P values for significance were calculated using the mean for each replicate and are displayed via SuperPlot, according to^42^ where the replicate mean is represented by a larger circle size than for subsample measurements. For p values > 0.05, not significant is indicated as “ns” and as statistically significant with * for p ≤ 0.05, ** for p ≤ 0.01, *** for p ≤ 0.001, **** for p ≤ 0.0001. Error is reported as plus or minus standard error of the mean, unless otherwise indicated.

### Plasmid Generation

Mutations in profilin were generated from EGFP-PFN1 (Addgene: 56438) or from synthesized β-actin-profilin constructs using site-directed mutagenesis (Q5 New England Biolabs, H133S and R88E) with primers in Table 1. Mutations in β-actin to render it polymerization-incompetent were generated from mCherry-β-actin (Addgene: 54967) or eGFP-β-actin (21948) or synthesized β-actin-profilin constructs. β-actin-profilin constructs were created using the IDT gblock synthetic gene construction service and cloned into pEGFP-C2 or pmCherry-C1 (Clontech, Mountainview, CA) using HindIII, BamHI. Codon optimization was performed within the manufacturer’s specifications limiting nucleotide repeats and codons were optimized for mus musculus. Sequences constructed as gblocks include: β-actinGS_16_profilin1_WT_, β-actinGS_30_profilin1_WT_, β-actinGS_30_profilin2_WT_, WAVE1PRD_WT_,WAVE1PRD_LOW_, and WAVE1PRD_MUT_. Mutagenesis was confirmed twice by sequencing (Azenta, Primordium) using both primer-based sequencing and whole-plasmid primer-less sequencing to ensure no off-target effects. See primer table 1.

### Microscopy and Image Analysis

Most imaging was performed at the UCSF Center for Advanced Light Microscopy (CALM). All FRAP experiments were performed using spinning disk microscopy with a TIRF Apo 60X (NA 1.49) objective on a GE OMX-SR microscope (inverted) with 3 PCO 15bit pco.edge sCMOS cameras. Cells were maintained at 37°C and 5% CO_2_ during imaging using an Okolab stage top incubator and controller. FRAP spot size was 1µm as measured from the surface of the coverslip. Analysis of fluorescence microscopy data was performed using FIJI^43^ or custom Matlab code as indicated. For correlation analysis, custom Matlab code was developed to identify and track fluorescence intensity with the dynamics of the leading edge of the cell. Time-lapse images consist of 100 frames of each fluorophore, where frames were acquired at a rate of 1 frame/second. The β-actin channel was used to identify the cell edge and a dynamic region of interest was created as input for an adaptive kymograph. Channels were first intensity-corrected for photobleaching and background. Tracking and plotting parameters include: gaussian blurring to smooth images, thresholding to identify the cell edge, and normalization of channels to their maximum intensities for plotting. To normalize each image, the channel with the lesser mean intensity was multiplied by a factor to match the mean intensity of the corresponding channel, so that each cell had the same mean intensity. Each analysis produced one kymograph for each channel averaged over the same spatial dimension indicated at the leading edge. Intensity traces were aligned by computing line by line cross-correlation in the time dimension from Matlab’s internal cross-correlation function (x-corr). Linear regression was used to find the best-fit slope of the line through the origin as well as the correlation coefficient. Each line was normalized to a “perfect” correlation of β-actin with itself using two different fluorophores (Fig. S7). All code is available in a public repository (link).

For display of ratiometric imaging data, image calculations were performed on the whole image in ImageJ. Images were first background-subtracted. To normalize each image, the channel with the lesser mean intensity was multiplied by a factor to match the mean intensity of the corresponding channel, so that each cell had the same mean intensity after thresholding. Channel LUTs were then identically scaled. Ratio images were generated by dividing the normalized images, in this case, the actinGS_30_profilin_WT_ or actinGS_30_profilin_PBD_ channel by the β-actin channel.Fluorescence intensity comparisons were made by drawing lines perpendicular to the cell boundary. Measurements were focused on quantifying actin intensity in the lamellipod and care was taken to ensure that lines did not include filopodia (Fig. S6).

### Retrograde Flow

Cells co-transfected with β-actin-profilin constructs and β-actin alone were plated onto laminin-coated coverslips for 45 min and then imaged with TIRF microscopy at 1 frame/s. The β-actin channel was used for analysis after import into ImageJ. Retrograde flow was calculated by measuring the distance and time that fiduciary markers traveled in the kymograph and solving for rate. Three biological replicates were performed per condition, unless stated otherwise.

### Morphology and Principal Component Analysis

Cell outlines and morphology measurements were generated in FIJI and the following measurements were taken: Solidity: (A_c_/A_h_, where A_c_ and A_h_ are the areas of cell and convex hull) Circularity: 4π(area/perimeter^2^)

Principal component analysis and visualization was performed in R with the “FactoMineR” and “FactoExtra” (ver. 1.0.6) packages. Analysis of the principal components yields the following: both circularity and solidity contribute equally but negatively to principal component 1 while the difference between these two variables is captured by principal component 2. Groups with higher principal component 1 scores (for instance, B16 WAVE1/2 knockouts) have lower circularity and solidity values while groups with lower principal component 1 scores, such as B16F1 wildtype cells, have higher circularity and solidity values.

### Cell Culture

B16F1 (ATCC-CRL-6323) mouse melanoma cells were cultured in DMEM/F12 with 4.5 g/mL glucose and supplemented with 10% fetal bovine serum (FBS, Life Technologies Certified, US) and penicillin/streptomycin. Cells were maintained at 37°C and 5% CO2 and split every 2–3 days. Cells were routinely confirmed to be free of myocplasma using Mycostrip (Invivogen). B16F1 WAVE1/2 knockouts and control knockouts were a generous gift from Bruce Goode (Brandeis University) and were maintained identically to wildtype. Depletion of WAVE1 and WAVE2 was independently confirmed by western blot (Fig.. 9).

### Live Cell Imaging

The Neon Transfection System (Invitrogen) was used to introduce DNA constructs into cells using the 10 μL transfection kit. Briefly, cells were grown to a confluency of 70%–80%, trypsinized and pelleted by centrifugation (160g). The pellet was rinsed with DPBS and resuspended in a minimal amount of buffer R (Invitrogen) with a total of 1μg of DNA or 0.1 μg/ μL of cell suspension. Cells transfected with DNA constructs were given 14-18 h after transfection before further experimental procedures were performed. Prior to imaging, transfection efficiency was visually confirmed on an EVOS FL digital inverted microscope objective (Life Technologies) equipped with a 10X Plan achromat 0.25 N.A. objective. Experiments were not performed on cells with low transfection efficiency (∼<50%).

Live cell imaging experiments were performed on 35mm Mattek dishes with 1.5 coverslips and 14mm glass diameter (P35G-1.5-14-C). Prior to day of imaging, surfaces were prepared by plasma cleaning for 5 minutes and coating with 50 μg/μL mouse laminin (Sigma Cat# L2020) diluted in phosphate-buffered saline. Glass was incubated with laminin overnight at 4°C. On the day of imaging, laminin-coated glass was rinsed with PBS. B16F1 cells were trypsinized and plated on laminin-coated glass with imaging medium (DMEM/F12 medium + 10% FBS + 50 mM HEPES, no phenol red) and were given 30 min to adhere.

### Immunofluorescence

Glass surfaces were prepared as previously described and cells were given 45 minutes to adhere prior to fixation with 4% electron microscopy grade paraformaldehyde (Electron Microscopy Sciences) for 10 min at room temperature. Cells were then permeabilized for 3 min with 0.1% Tween-20 and washed three times with PBS, then blocked for 20 minutes in 5% bovine serum albumin. Cells were then stained overnight at 4°C with primary antibodies diluted in 1% bovine serum albumin in PBS. They were then washed twice with PBS for 5 min and incubated with secondary antibodies (diluted 1:1000) for 1 h at room temperature in PBS. Actin filaments were stained with Alexa Fluor 488 phalloidin or Alexa Fluor 568 phalloidin (diluted 1:100, Life Technologies) for 30 min at room temperature in immunofluorescence staining buffer. Cells were washed three times with PBS before mounting with Prolong Diamond (Life Technologies). The following antibodies/dilutions were used: Fascin (55k-2) mouse (Cell Signaling Technology, 54545S, 1:200), Vasp (9A2) Rabbit (Cell Signaling Technology, 3132T, 1:400). Anti-mouse IgG 647 and anti-rabbit IgG 568 (Life Technologies) were used at 1:1000 dilution.

### Western blots

Adherent cells were harvested with a cell scraper in RIPA buffer with cOmplete EDTA-free Protease Inhibitor Cocktail Roche (Millipore Sigma). Whole cell lysates were prepared by membrane disruption using repeated passage through a 27 gauge needle. Protein content was then assessed with Pierce BCA Protein Assay Kit (Thermo Fisher) and diluted in SDS buffer stained with Orange G (40% glycerol, 6% SDS, 300 mM Tris HCl, pH 6.8). 10μg samples were evenly loaded on an SDS-PAGE gel (Bio-RAD 4%–20% Mini-Protean TGX Gels). Protein was transferred to a PVDF membrane (0.2 micron, Immobilon) and blocked in 5% Bovine Serum Albumin (BSA) (Sigma-Aldrich) for 30 minutes. All antibodies were diluted in 5% BSA and 0.1% Tween-20 (Fisher Scientific). Primary antibodies were incubated at 4°C overnight and secondary antibodies (Li-Cor; Abcam) were incubated for 2 h at room temperature. WAVE1,2 isoforms or GAPDH from whole cell lysate were detected with Li-Cor fluorescent antibodies on an Odyssey detection system (Li-Cor) WesternSure Pre-Stained Chemiluminescent Protein Ladder (Li-Cor) was used as a molecular weight marker. The following antibodies/dilutions were used: WASF1/WAVE1 Antibody (S03-5K1, Novus Biologicals, 1:1000), WAVE-2 (D2C8) XP® Rabbit mAb (Cell Signaling Technology, 3656T), GAPDH Mouse Monoclonal Antibody [D4C6R] (Cell Signaling Technology, 97166T).

### Molecular Dynamics Simulations and Analysis

For molecular simulations, we started with the actin-profilin co-complex (PDB 1HLU)^44^. Missing atoms/residue were rebuilt resulting in the unlinked complex used as a control. The cross-linked complexes were built using MoMA^45^ with sequences of G4SG4S for the 10-AA cross-link and G4SG4SG5S for the 16-AA cross-link. Each of these three systems were solvated using TIP3P water with 10 Å padding. Na+ and Cl-were added to neutralize the system and give an ionic strength of 50 mM. NAMD 3.0^46^ was used to perform the simulations using the CHARMM36 forcefield^47^. Following minimization, the systems were heated and equilibrated at a temperature of 300 K and 1 atm of pressure (NpT conditions). Simulations were performed with 2 fs timesteps by fixing hydrogens. We employed Particle Mesh Ewald^48^ for long-range electrostatics and used a 10 Å cut-off and 8.5 Å switch distance for van der Waals interactions. After equilibration for 100 ns, each system had two independent simulations of 0.5 µs each, resulting in 1 µs for each and 3 µs overall.

All analysis was performed using Bio3d^49^ in R. From visualization of the simulations, it was clear that the shorter, 10-AA linker caused the actin-profilin complex to be distorted. To quantify this, we performed PCA on the Cα atoms for the residues that constitute up the binding interface between actin and profilin. Projecting all the simulations onto the first PC vector of the principal component space showed that the 16-AA linker had minimal effects on the actin-profilin complex while the shorter linker caused a significant conformational shift.

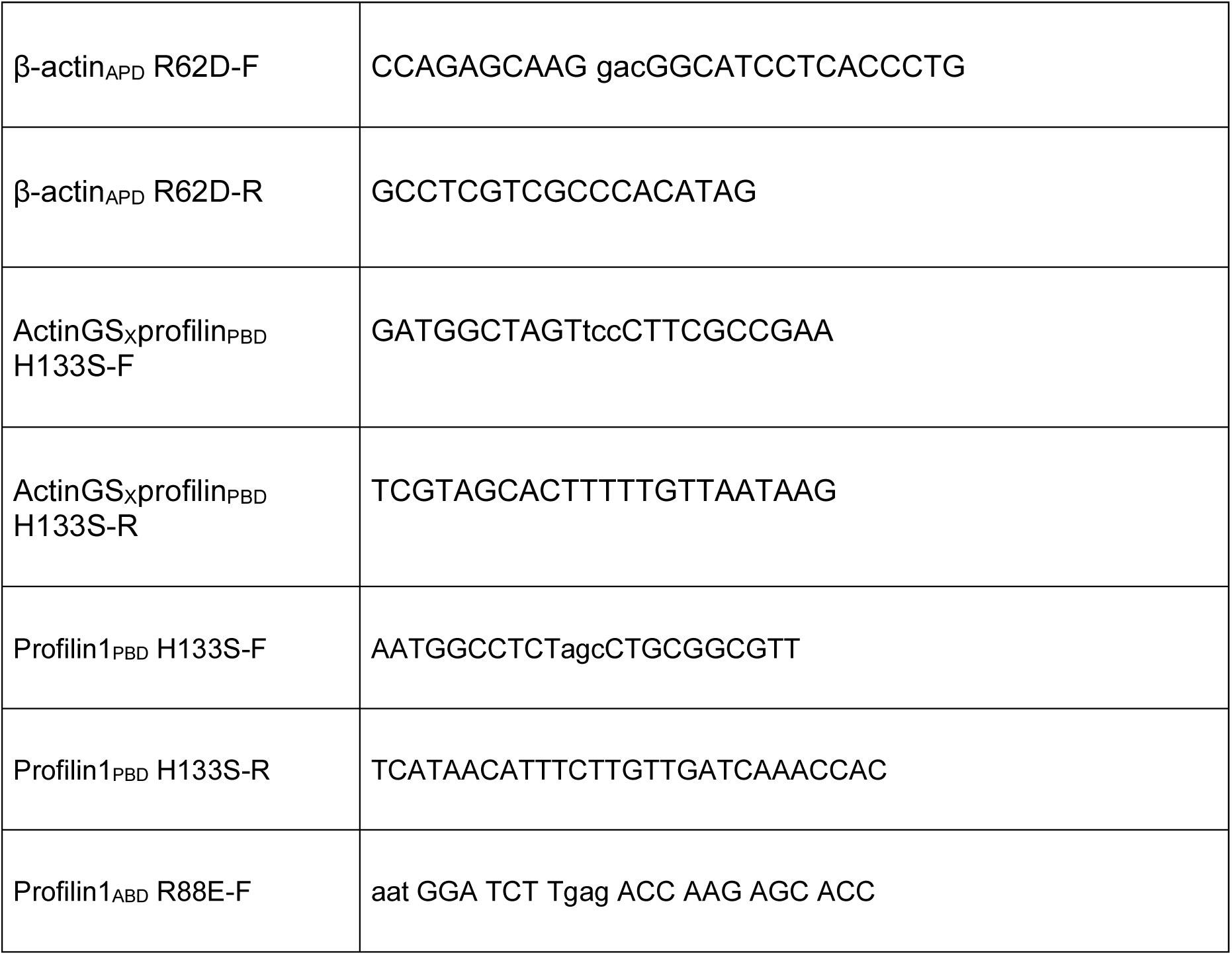
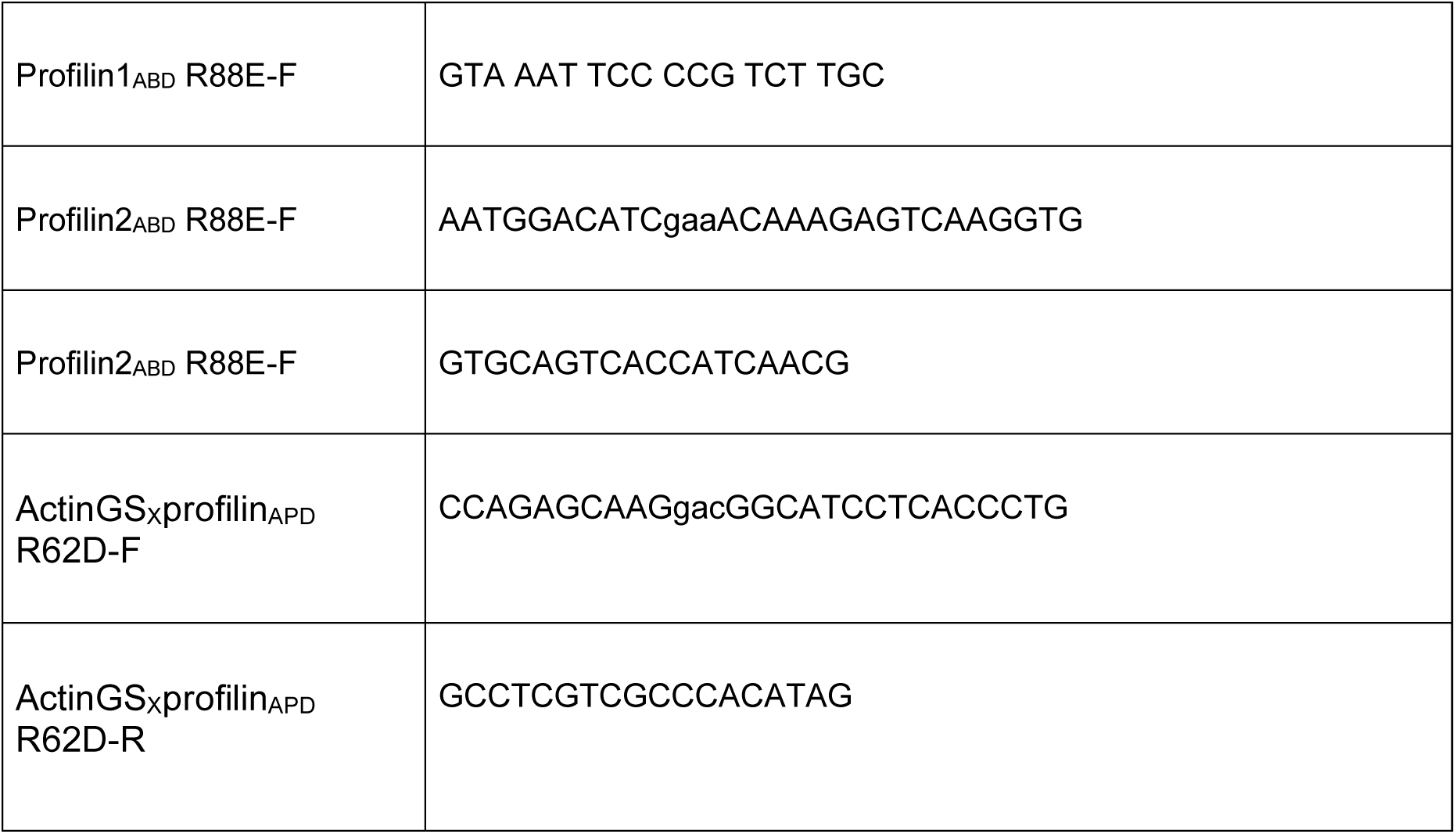
Primer Table

## Acknowledgments

We would like to thank members of the Mullins lab for reagents, experimental advice and feedback on the manuscript, with extra special thanks to Sam Lord and Natalie Petek. We would also like to thank Beryl Rappaport for careful manuscript review and perspective. Additional thanks to the UCSF Center for Advanced Light Microscopy (CALM) for assistance with imaging, particularly Kari Harrington for help with performing RICS and Michelle Digman and Enrico Gratton (University of California, Irvine) for their help with RICS analysis. We would lastly like to thank the Goode lab for sharing their B16F1 WAVE1/2 knockouts with us. This work was supported by the National Institute of General Medical Sciences of the National Institutes of Health (R35-GM118119 to R.D.M.), by the Howard Hughes Medical Institute Investigator program (R.D.M.)

## References

1. Drenckhahn, D. & Pollard, T. D. Elongation of actin filaments is a diffusion-limited reaction at the barbed end and is accelerated by inert macromolecules. J. Biol. Chem. 261, 12754– 12758 (1986).

2. Mullins, R. D., Bieling, P. & Fletcher, D. A. From solution to surface to filament: actin flux into branched networks. Biophys. Rev. 10, 1537–1551 (2018).

3. Fujiwara, I., Vavylonis, D. & Pollard, T. D. Polymerization kinetics of ADP-and ADP-Pi-actin determined by fluorescence microscopy. Proc. Natl. Acad. Sci. 104, 8827–8832 (2007).

4. Bieling, P. et al. WH2 and proline-rich domains of WASP-family proteins collaborate to accelerate actin filament elongation. EMBO J. 37, 102–121 (2018).

5. Mullins, R. D. & Skruber, K. Actin filament assembly driven by distributive polymerases clustered on membrane surfaces. 2024.11.26.625540 Preprint at 10.1101/2024.11.26.625540 (2024).

6. Bisi, S. et al. Membrane and actin dynamics interplay at lamellipodia leading edge. Curr. Opin. Cell Biol. 25, 565–573 (2013).

7. Mooren, O. L., Galletta, B. J. & Cooper, J. A. Roles for Actin Assembly in Endocytosis. Annu. Rev. Biochem. 81, 661–686 (2012).

8. Insall, R. H. & Machesky, L. M. Actin dynamics at the leading edge: from simple machinery to complex networks. Dev. Cell 17, 310–322 (2009).

9. Efimova, N. & Svitkina, T. M. Branched actin networks push against each other at adherens junctions to maintain cell-cell adhesion. J. Cell Biol. 217, 1827–1845 (2018).

10. Richardson, B. E., Beckett, K., Nowak, S. J. & Baylies, M. K. SCAR/WAVE and Arp2/3 are crucial for cytoskeletal remodeling at the site of myoblast fusion. Dev. Camb. Engl. 134, 4357–4367 (2007).

11. Clark, A. G. et al. Integration of single and multicellular wound responses. Curr. Biol. CB 19, 1389–1395 (2009).

12. Kast, D. J. & Dominguez, R. WHAMM links actin assembly via the Arp2/3 complex to autophagy. Autophagy 11, 1702–1704 (2015).

13. Zalevsky, J., Lempert, L., Kranitz, H. & Mullins, R. D. Different WASP family proteins stimulate different Arp2/3 complex-dependent actin-nucleating activities. Curr. Biol. CB 11, 1903–1913 (2001).

14. Mullins, R. D. & Pollard, T. D. Structure and function of the Arp2/3 complex. Curr. Opin. Struct. Biol. 9, 244–249 (1999).

15. Machesky, L. M. & Gould, K. L. The Arp2/3 complex: a multifunctional actin organizer. Curr. Opin. Cell Biol. 11, 117–121 (1999).

16. Machesky, L. M. & Insall, R. H. Scar1 and the related Wiskott-Aldrich syndrome protein, WASP, regulate the actin cytoskeleton through the Arp2/3 complex. Curr. Biol. CB 8, 1347–1356 (1998).

17. Funk, J. et al. Profilin and formin constitute a pacemaker system for robust actin filament growth. eLife 8, e50963 (2019).

18. Chereau, D. & Dominguez, R. Understanding the role of the G-actin-binding domain of Ena/VASP in actin assembly. J. Struct. Biol. 155, 195–201 (2006).

19. Bestul, A. J. et al. Fission yeast profilin is tailored to facilitate actin assembly by the cytokinesis formin Cdc12. Mol. Biol. Cell 26, 283–293 (2015).

20. Pollard, T. D. & Cooper, J. A. Quantitative analysis of the effect of Acanthamoeba profilin on actin filament nucleation and elongation. Biochemistry 23, 6631–6641 (1984).

21. Lu, J. & Pollard, T. D. Profilin Binding to Poly-l-Proline and Actin Monomers along with Ability to Catalyze Actin Nucleotide Exchange Is Required for Viability of Fission Yeast. Mol. Biol. Cell 12, 1161–1175 (2001).

22. Gutsche-Perelroizen, I., Lepault, J., Ott, A. & Carlier, M.-F. Filament Assembly from Profilin-Actin*. J. Biol. Chem. 274, 6234–6243 (1999).

23. Nyman, T., Page, R., Schutt, C. E., Karlsson, R. & Lindberg, U. A cross-linked profilin-actin heterodimer interferes with elongation at the fast-growing end of F-actin. J. Biol. Chem. 277, 15828–15833 (2002).

24. Hahne, P., Sechi, A., Benesch, S. & Small, J. V. Scar/WAVE is localised at the tips of protruding lamellipodia in living cells. FEBS Lett. 492, 215–220 (2001).

25. Smith, M. B., Kiuchi, T., Watanabe, N. & Vavylonis, D. Distributed Actin Turnover in the Lamellipodium and FRAP Kinetics. Biophys. J. 104, 247–257 (2013).

26. Wang, Y. L. Exchange of actin subunits at the leading edge of living fibroblasts: possible role of treadmilling. J. Cell Biol. 101, 597–602 (1985).

27. van Rosmalen, M., Krom, M. & Merkx, M. Tuning the Flexibility of Glycine-Serine Linkers To Allow Rational Design of Multidomain Proteins. Biochemistry 56, 6565–6574 (2017).

28. Schutt, C. E., Myslik, J. C., Rozycki, M. D., Goonesekere, N. C. W. & Lindberg, U. The structure of crystalline profilin–β-actin. Nature 365, 810–816 (1993).

29. Ezezika, O. et al. Incompatibility with formin Cdc12p prevents human profilin from substituting for fission yeast profilin: insights from crystal structures of fission yeast profilin. J. Biol. Chem. 284, 2088–2097 (2009).

30. Cheng, K. W. & Mullins, R. D. Initiation and disassembly of filopodia tip complexes containing VASP and lamellipodin. Mol. Biol. Cell 31, 2021–2034 (2020).

31. Mouneimne, G. et al. Differential remodeling of actin cytoskeleton architecture by profilin isoforms leads to distinct effects on cell migration and invasion. Cancer Cell 22, 615–630 (2012).

32. Dominguez, R. The WH2 Domain and Actin Nucleation: Necessary but Insufficient. Trends Biochem. Sci. 41, 478–490 (2016).

33. Skruber, K. et al. Arp2/3 and Mena/VASP Require Profilin 1 for Actin Network Assembly at the Leading Edge. Curr. Biol. CB 30, 2651–2664.e5 (2020).

34. Yang, C. et al. Novel roles of formin mDia2 in lamellipodia and filopodia formation in motile cells. PLoS Biol. 5, e317 (2007).

35. Buracco, S. et al. Scar/WAVE drives actin protrusions independently of its VCA domain using proline-rich domains. Curr. Biol. CB 34, 4436–4451.e9 (2024).

36. Tang, Q. et al. WAVE1 and WAVE2 have distinct and overlapping roles in controlling actin assembly at the leading edge. Mol. Biol. Cell 31, 2168–2178 (2020).

37. Belin, B. J., Goins, L. M. & Mullins, R. D. Comparative analysis of tools for live cell imaging of actin network architecture. Bioarchitecture 4, 189–202 (2014).

38. Jacquemet, G., Hamidi, H. & Ivaska, J. Filopodia Quantification Using FiloQuant. Methods Mol. Biol. Clifton NJ 2040, 359–373 (2019).

39. Huang, W. Y. C., Boxer, S. G. & Ferrell, J. E. Membrane localization accelerates association under conditions relevant to cellular signaling. Proc. Natl. Acad. Sci. 121, e2319491121 (2024).

40. Padrick, S. B., Doolittle, L. K., Brautigam, C. A., King, D. S. & Rosen, M. K. Arp2/3 complex is bound and activated by two WASP proteins. Proc. Natl. Acad. Sci. 108, E472– E479 (2011).

41. Co, C., Wong, D. T., Gierke, S., Chang, V. & Taunton, J. Mechanism of Actin Network Attachment to Moving Membranes. Cell 128, 901–913 (2007).

42. Lord, S. J., Velle, K. B., Mullins, R. D. & Fritz-Laylin, L. K. SuperPlots: Communicating reproducibility and variability in cell biology. J. Cell Biol. 219, e202001064 (2020).

43. Schindelin, J., et al. Fiji: an open-source platform for biological-image analysis. Nat. Methods 9, 676–682 (2012).

44. Chik, J. K., Lindberg, U. & Schutt, C. E. The Structure of an Open State of β-Actin at 2.65 Å Resolution. J. Mol. Biol. 263, 607–623 (1996).

45. Barozet, A. et al. MoMA-LoopSampler: a web server to exhaustively sample protein loop conformations. Bioinformatics 38, 552–553 (2022).

46. Phillips, J. C. et al. Scalable molecular dynamics on CPU and GPU architectures with NAMD. J. Chem. Phys. 153, 044130 (2020).

47. Huang, J. & MacKerell Jr, A. D. CHARMM36 all-atom additive protein force field: Validation based on comparison to NMR data. J. Comput. Chem. 34, 2135–2145 (2013).

48. Essmann, U. et al. A smooth particle mesh Ewald method. J. Chem. Phys. 103, 8577– 8593 (1995).

49. Grant, B. J., Rodrigues, A. P. C., ElSawy, K. M., McCammon, J. A. & Caves, L. S. D. Bio3d: an R package for the comparative analysis of protein structures. Bioinforma. Oxf. Engl. 22, 2695–2696 (2006).

