## Supplemental Figures and Data for "Membrane-associated polymerases deliver most of the actin subunits to a lamellipodial network"

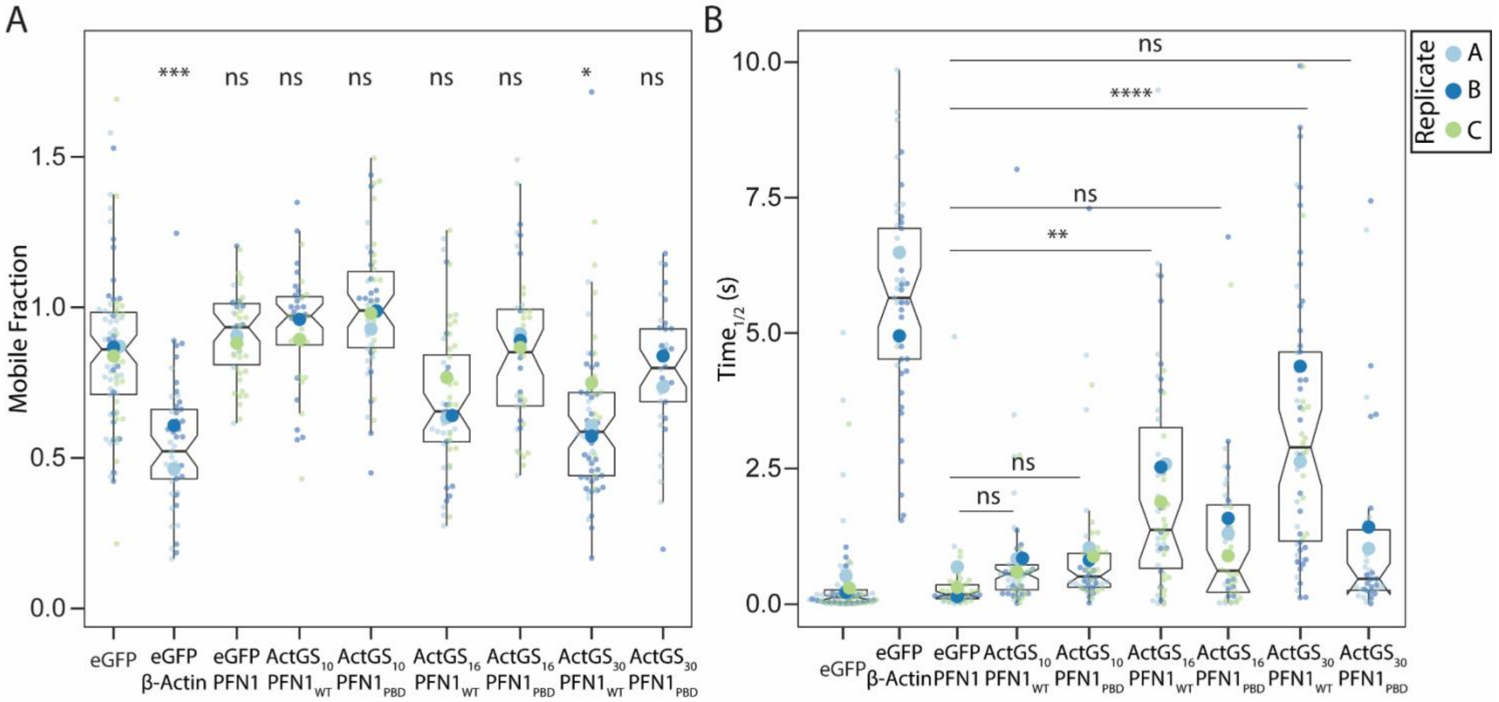


Fig. S1, related to Figure 1.

A. Quantification of mobile fraction of recovery curve after photobleaching per cell. B. Quantification of half-time of recovery curve after photobleaching per cell. Number of cells per experiment per round for both A and B are as follows: eGFP (37,23,22) β-actin$_{WT}$ (22,27), eGFP-profilin1 (11,9,26), actinGS$_{10}$profilin$_{WT}$ (23,18), actinGS$_{10}$profilin$_{PBD}$ (16,17,22), actinGS$_{16}$profilin$_{WT}$ (21,12,23), actinGS$_{16}$profilin$_{PBD}$(15,11,18), actinGS$_{30}$profilin$_{WT}$ (16,35,14), actinGS$_{30}$profilin$_{PBD}$ (19,19). One-way ANOVA and Tukey's multiple comparison tests are comparisons with eGFP for A and indicated groups for B.

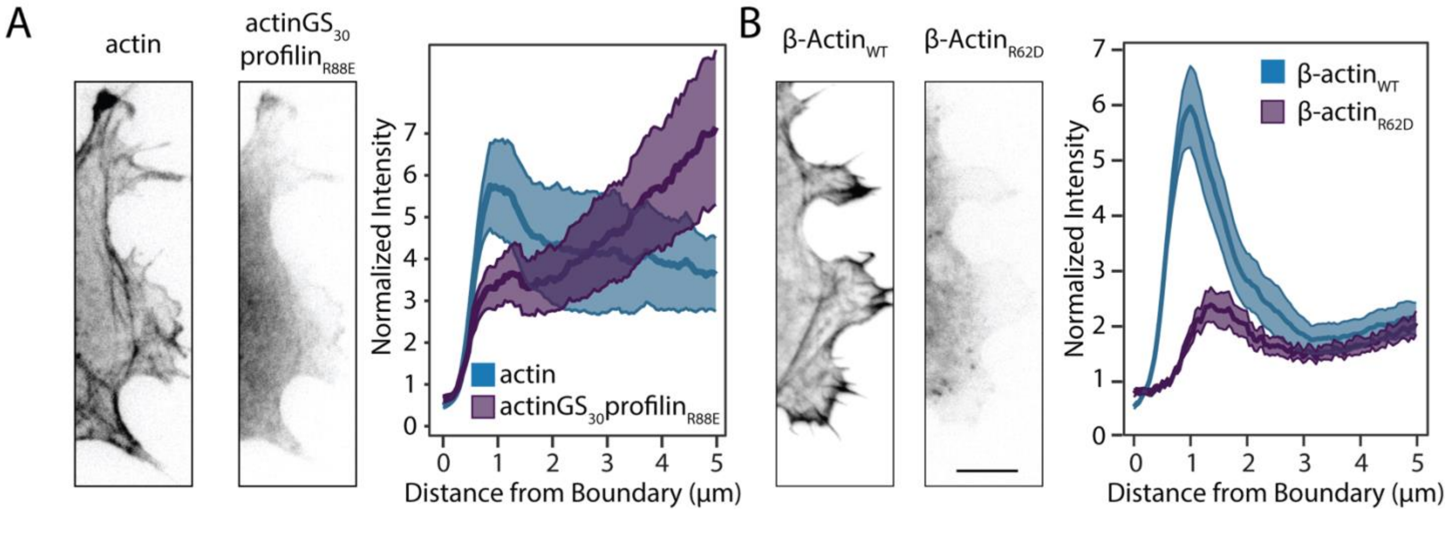


Fig S2. A. Representative images and averaged line scans of actin and actinGS$_{30}$profilin$_{R88E}$ fluorescence intensity at the leading edge of cells. Scale bar represents 5µm. Profilin$_{R88E}$ is a profilin mutant not competent to bind actin. B. Representative images and line scan analysis of β-actin and β-actin$_{R62D}$ fluorescence intensity at the leading edge of cells. β-actin$_{R62D}$ is a polymerization incompetent mutant of actin. The cell edge= 0 and bands depict 95% confidence intervals. All lines are normalized to the cell mean intensity. For all conditions, n = 100 lines drawn from 10 cells.

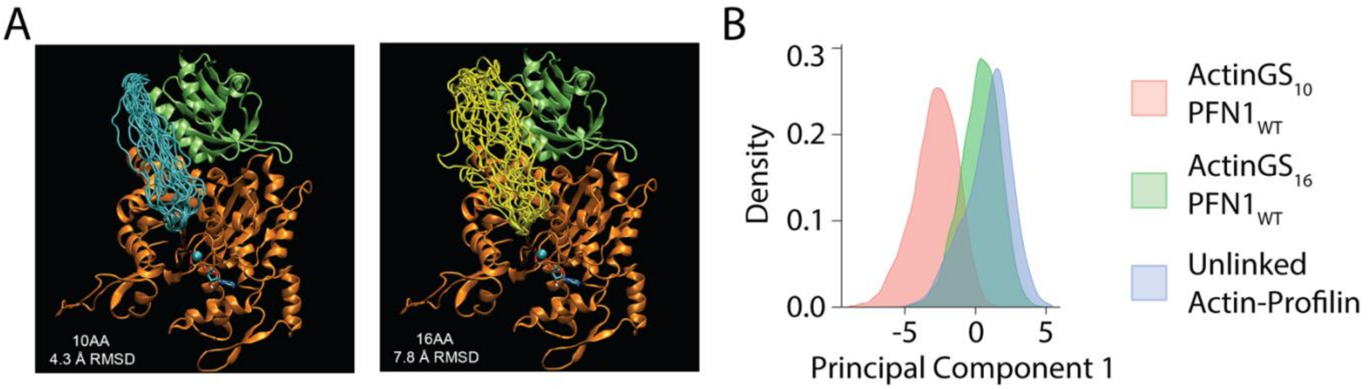


Fig. S3, related to Figure 1. A. Illustration of actin-profilin orientations for linked constructs using previously published structures (PDB:2BTF) with either 10 or 16 alternating glycine-serine residues. Principal component analysis comparing the mobility of actinGS$_{10}$profilin$_{WT}$ and actinGS$_{16}$profilin$_{WT}$ with unlinked actin-profilin.

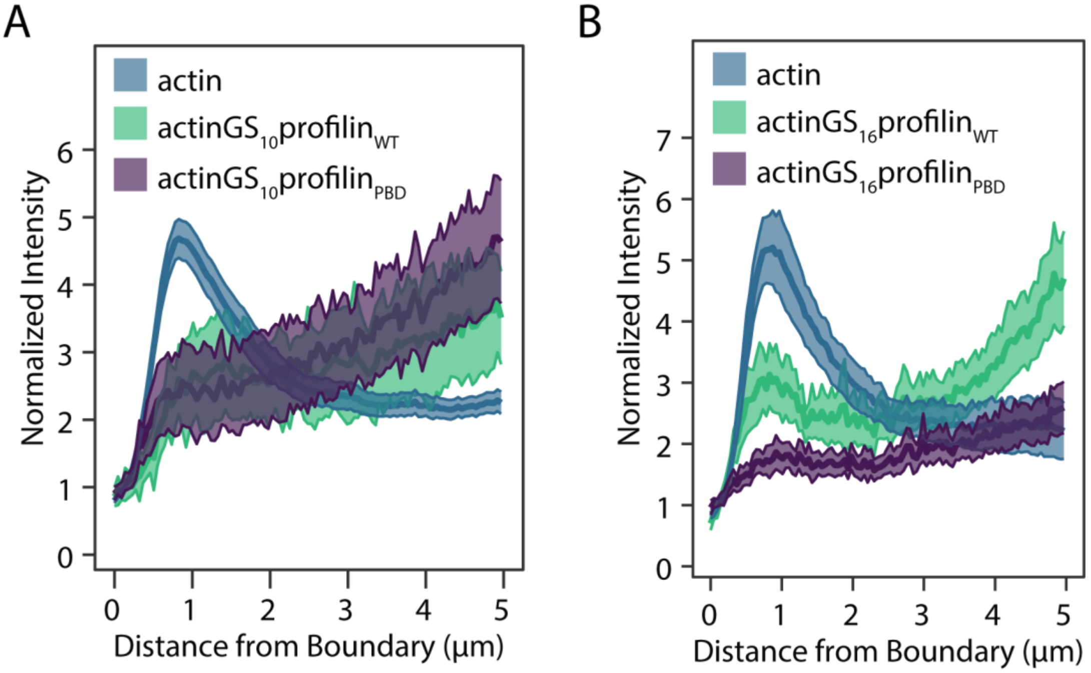


Fig. S4, related to Figure 1. Averaged linescans of (A) actinGS$_{10}$profilin$_{WT}$ and proline-binding deficient actinGS$_{10}$profilin$_{PBD}$ (B) actinGS$_{16}$profilin$_{WT}$ and proline-binding deficient actinGS$_{16}$profilin$_{PBD}$ fluorescence intensity at the leading edge of cells in comparison to actin. The cell edge = 0 and bands depict 95% confidence intervals. For all conditions, n = 100 lines drawn from 10 cells. All lines are normalized to the cell mean intensity.

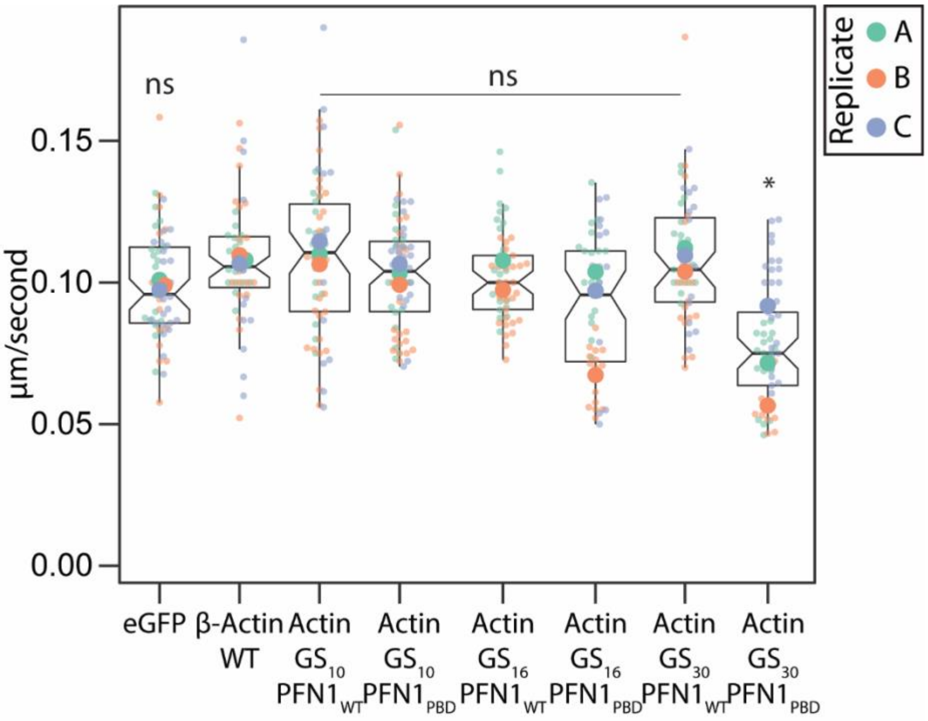


Fig. S5, related to Figure 3. Quantification of retrograde flow of mCherry-β-actin with co-transfection of indicated construct. Number of measurements per experiment per round where 1 kymograph was generated per cell are as follows: eGFP (20,16,25), $actin_{WT}$ (20,20,15), $actinGS_{10}profilin_{WT}$ (20,31,15), $actinGS_{10}profilin_{PBD}$ (20,25,25), $actinGS_{16}profilin_{WT}$ (23,30), $actinGS_{16}profilin_{PBD}$(20,15,16), $actinGS_{30}profilin_{WT}$ (20,20,15), $actinGS_{30}profilin_{PBD}$ (20,11,53). One-way ANOVA and Tukey's multiple comparison tests are comparisons with actin.

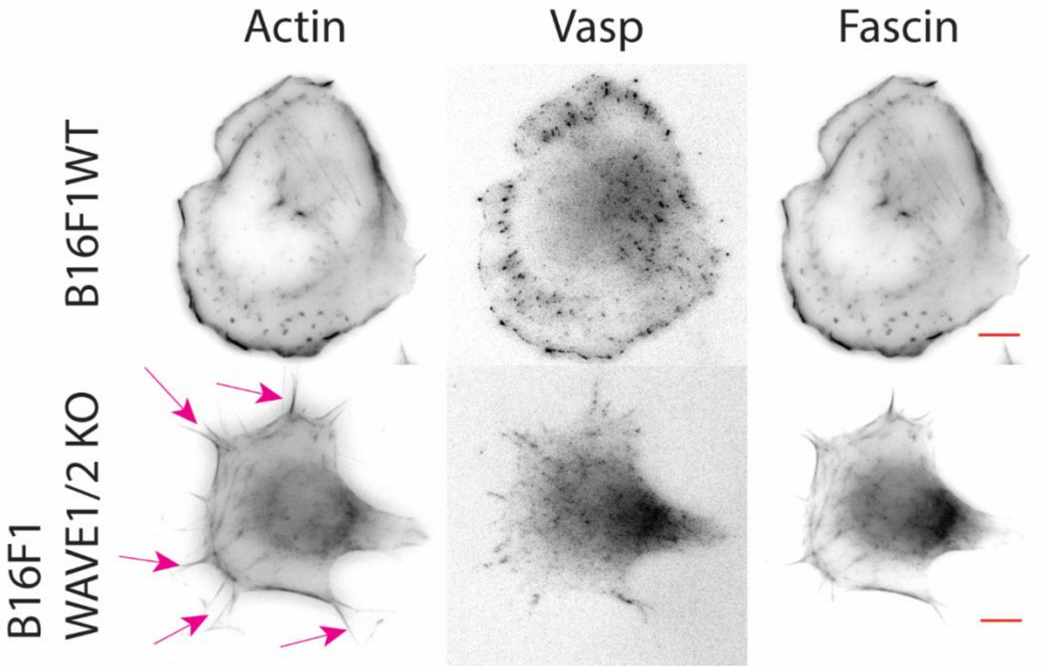


Fig. S6, related to Figure 4. Representative images of Vasp and Fascin immunolabeling and phalloidin647 staining in B16F1 control and B16F1 WAVE1/2 KO cells. Staining shows that long linear filopodia-like structures in B16F1 WAVE1/2 are positive for filopodial markers. Arrowheads point to filopodia. Scale bar represents 10µm.

A

| Groups | Dim. 1 | Dim. 2 |
| --- | --- | --- |
| B16F1$_{WT}$ | -0.73 | -0.11 |
| WAVE1/2$_{KO}$ | 1.32 | 0.23 |
| KO+ WAVE1$_{MUT}$ | 0.33 | -0.12 |
| KO+ WAVE1$_{WT}$ | -0.90 | 0.10 |

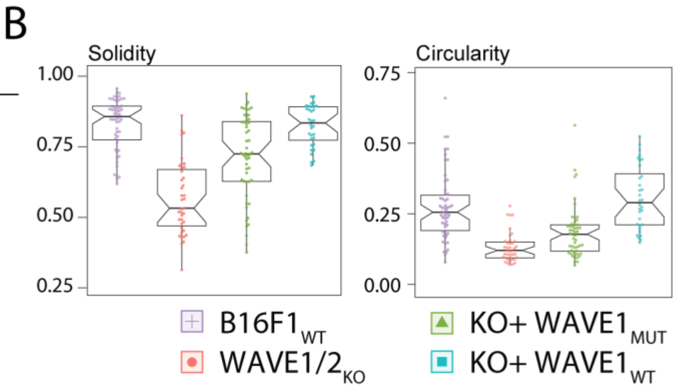


Fig. S7, related to Figure 4. Principal component analysis of B16F1 control and B16F1 WAVE1/2 KO cell shape. A. Mean coordinates of individual samples for each group per dimension. C. Cell shape data from 54 B16F1 WT, 60 B16F1 WAVE1/2 knockout, and 52,60  B16F1 WAVE1/2 knockout cells expressing WAVE1PRD$_{MUT}$ or WAVE1PRD$_{WT}$, respectively.

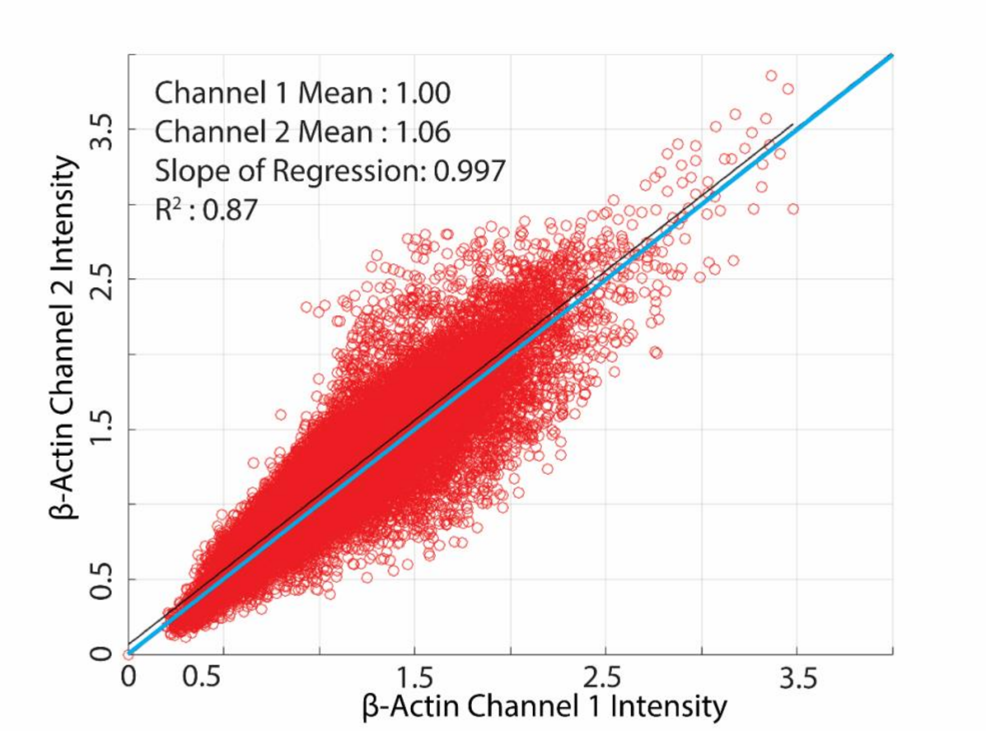


Fig. S8, Related to Methods. Linear correlation lines for intensity plots are fit and normalized to an ideal “perfect” correlation of actin with itself using two different fluorophores. Each channel is thus divided by the mean of the actin and the reference actin correlation line is displayed as a thick black line through the origin. The line is highlighted here in cyan for clarity.

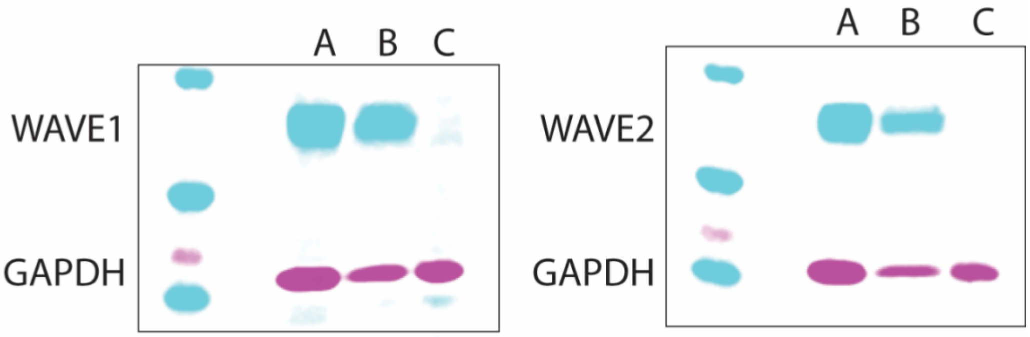


Fig. S9, related to Methods. Representative western blots of B16F1 WAVE1/2 knockout cells with GAPDH as a loading control. Full validation is in (Tang reference). From left to right: (A) B16F1 wild-type cells (B) Control knockout cells generated in (C) WAVE1/2 knockout cells
